# Morphometric Characterization and Variation of Somali Dromedary Camels (*Camelus dromedarius*) in a selected district of Benadir Region

**DOI:** 10.64898/2026.04.12.718019

**Authors:** Hassan Hussein Abdi, Alper Yilmaz, Abdirahman Barre, Jesse Faez Firdaus Abdullah, Mohamed Abdullahi Mohamud

**Author notes:** **Corresponding author:** Dr. Abdirahman Barre, Department of Veterinary Clinical Studies, Faculty of Veterinary Medicine, University Putra Malaysia, 43400 Serdang, Selangor, Malaysia, /.

## Abstract

Camels are one of the hardy animal species known for their high adaptability to extreme climatic conditions. However, scientific studies on this important animal species are limited. This study aims to investigate the statistical relationships between some body measurements of dromedary camels with different characteristics raised in the Mogadishu, Somalia. The research was carried out on a total of 248 camels (102 males, 146 females) of different age groups, belonging to the SiifDacar, Hoor, and Eyddimo types, raised in camel farms in Dharkenley, Dayniile, and Hodan districts of Mogadishu. Body measurements were taken manually using a standard measuring tape, and the collected data were subjected to statistical analysis. As a result of the analyses, significant differences were found between the camels named with the local language as mentioned above in terms of withers height, rump height, body length, and shoulder width in female camels. In male camels, it was determined that there was a statistically significant difference among the camels named in terms of rump width. When evaluated specifically for female camels, it was found that the camels of the SiifDacar had higher withers and rump height compared to the camels of the Hoor, and were also longer than the other two different camels. Although no notable difference in shoulder width was found between the Hoor and Eyddimo camels, Siif Dacar females exhibited broader shoulders than both. In male camels, Siif Dacar had a significantly wider rump compared to those of the Hoor camels. Furthermore, the study identified varying degrees of statistically significant correlations among different body measurements in the camels. Overall, it was observed that female camels of the Siif Dacar were larger in size compared to those of the Hoor and Eyddimo camels, and that male camels generally exhibited greater body dimensions than females. These findings contribute to the registration of the different-named camel types in Somalia as separate breeds and to a better understanding of the morphological characteristics of camels.

## Introduction

Somalia is located in the Horn of Africa, between the latitudes of 5.1521° N and the longitudes of 46.1996° E, and it has a surface area of approximately 638,000 square kilometers (Abdulraheem *et al*., 2024). The capital city of Somalia is Mogadishu, and the country generally exhibits arid and semi-arid climatic characteristics. These climatic conditions provide Somalia with great potential for nomadic livestock farming (AlAsmari and Mir, 2026). The livestock sector makes a significant contribution to the country’s gross domestic product (GDP) and constitutes the primary source of livelihood. The main livestock species raised in the country include sheep, goats, camels, and cattle (Muigai et al., 2016). Additionally, Somalia is influenced by air currents originating from the Indian Ocean and by the northeastern winds coming from Arabia and Asia, and it has four main seasons. Among these, the Gu season (April-June) receives the heaviest rainfall, while the Deyr season (October-December) is characterized by lighter and more sporadic precipitation (Baumann, 2014). These climatic conditions, particularly, increase the importance of camels in the livestock sector (Fadlelmoula *et al*., 2015).

Camels are utilized in a wide range of sectors, including dairy and meat production, wool harvesting, transportation, agricultural labor, sports activities, and tourism (Fares, 2025). Globally, it is estimated that the camel population comprises approximately 31.1 million dromedary camels (Camelus dromedarius) and 3.7 million Bactrian camels (Camelus bactrianus). A substantial proportion of this population, nearly 60%, is concentrated in Northeastern African countries, particularly Somalia, Sudan, Ethiopia, and Kenya (FAO, 2017). Beyond their economic significance, camels hold an essential position within the cultural and social practices of the region, being symbolically and practically integrated into important life events such as marriages, funerals, and religious ceremonies (Camasepro, 2012).

The main types of camels raised in Somalia are named Siif Dacar, Eyddimo, and Hoor. The Siif Dacar is predominantly found in the southern parts of the country, especially in the Middle Shabelle Valley and the Central Coastal areas (Alhajeri *et al*. 2019). These camels are also raised by Somali communities residing in Kenya and are typically characterized by their white or sandy yellow coat color (Farah et al., 2014). The Eyddimo camels are located primarily in the borderlands between Somalia and Ethiopia, as well as along the hot coastal regions of Djibouti. Noted for their relatively small body size, they have adapted to severe environmental conditions and are distinguished by a reddish coat color (Hussein *et al*., 2019).

Among these, the Hoor camels are recognized as the most widely distributed camels in Somalia, with their primary habitat spanning regions such as Nogal, Bakool, Gedo, and the Central Shabelle Valley. As can be understood from here, camel types with different names have been raised in different regions for many years and thus have differentiated morphologically (Alhajeri *et al*., 2021).

The scientific determination of the body characteristics of animals holds considerable significance for livestock breeding practices and selection programs. Regular recording of body measurements provides valuable insights into the animals’ adaptation to ecological conditions, nutritional status, and general health. Furthermore, these measurements play a crucial role in estimating the commercial value of livestock (Barre, 2023).

The present study aims to identify specific body measurements of various Somali dromedary camel types in Mogadishu, with the intention of revealing the genetic and morphological diversity within the region and establishing a foundational dataset for future research. The findings obtained from this research may contribute to breed registration efforts and productivity analyses, while also offering critical data for the improvement of camel breeding practices, the development of selection programs, and the formulation of animal husbandry strategies specific to the region (Barre *et al*., 2025).

Within this context, the study compares the body measurements of different camel types, examines the influence of sex on these measurements, and evaluates the statistical relationships among the recorded parameters. Determining body measurements is essential for the identification of phenotypic traits in animals (Çağlı, 2019). Understanding how these traits vary among different breeds is vital for preserving breed characteristics and for designing appropriate breeding and improvement programs. Additionally, the data generated from this study provides essential information for the sustainable management of livestock production in the region and for developing more efficient production systems (Barre *et al*., 2023).

In this context, the relationships between the body parameters measured in the study were analyzed, contributing to the understanding of growth and development processes and evaluating whether body measurements are characteristic of the specified types. Such data may help to optimize feeding and management strategies and allow breeders to make more informed choices (Eisa, 2011).

## Materials and Methods

This study was conducted across five distinct camel farms located in the Dharkenley, Dayniile, and Hodan districts of Mogadishu, Somalia. A total of 248 dromedary camels, aged six years and older, representing three different Somali dromedary camel types, Siif Dacar, Hoor, and Eyddimo, were examined. The sample consisted of 102 males and 146 females. The animals evaluated in the study are summarized in Table 1 in terms of sex, age, and reported types.

**Table 1.** Distribution of Camels Included in the Study by Age, Sex, and Type.

| **Camel Type** | **Sex** | **6–9 years** | **10–13 years** | **14 years and above** | **Total** |
| --- | --- | --- | --- | --- | --- |
| SiifDacar | Female | 19 | 10 | 20 | 49 |
|  | Male | 7 | 17 | 9 | 33 |
| Hoor | Female | 26 | 16 | 15 | 57 |
|  | Male | 12 | 14 | 12 | 38 |
| Eyddimo | Female | 19 | 9 | 12 | 40 |
|  | Male | 7 | 16 | 8 | 31 |
| **Total** |  | **90** | **82** | **76** | **248** |

As presented in Table 1, the distribution of the camels analyzed in this study was structured with attention to maintaining balance across age, sex, and type. Of the total 248 animals examined, 146 were females and 102 were males, with representation across distinct age categories.

The number of Siif Dacar type included a total of 82 camels, comprising 49 females and 33 males. Among these camels, the largest number of animals are in the 14-year-old and above age group (n=29), while the number of camels in the 6-9 age group is 26. The Hoor camels were the most represented in the study, with 95 camels in total—57 females and 38 males. An analysis of age groups revealed that the majority of Hoor camels (n=38) were in the 6–9-year age range, while the 14+ age group had the fewest animals (n=27). A total of 71 camels from the Eyddimo type were evaluated, of which 40 were females and 31 were males. In this group, 26 camels fell within the 6–9-year age range, 25 were aged between 10–13 years, and 20 were in the 14 years and older category.

Overall, the camels assessed in this study exhibited a relatively balanced distribution across age groups. Nevertheless, some discrepancies in the number of individuals within specific age and sex subgroups were observed. For instance, in all three types, the number of female camels exceeded that of males. Additionally, there were 90 camels in the 6–9-year group, 82 in the 10–13-year group, and 76 in the 14+ group. Despite these variations, the distribution was deemed adequate for comparative analysis of body measurements across different age and sex categories. Accordingly, statistical evaluations were conducted to examine variations in morphometric traits associated with age and sex within each camel type.

### Measurement of Body Dimensions

In this study, the body measurements of camels were taken using a measuring tape in accordance with standard procedures described in the literature (Çağlı 2019), and all measurements were recorded in centimeters (cm). The animals were measured while standing still in an upright position to ensure consistency. A total of eight morphometric traits were assessed: withers height, back height, rump height, body length, chest height, abdominal height, shoulder width, and seat width. Withers height was measured as the vertical distance from the highest point of the withers to the ground, while back height was defined as the vertical distance from the spinous process of the last thoracic vertebra to the ground. Rump height was measured from the highest point of the pelvic region (sacrum) to the ground, and body length was determined as the distance between the caput humeri and the tuber ischiadicum (ischial tuberosity). Chest height and abdominal height were recorded as the vertical distances from the ground to the lowest points of the chest and abdomen, respectively. Shoulder width was defined as the distance between the two humeral bones, and seat width as the distance between the two femoral bones. To minimize potential measurement errors due to animal movement, assistance from animal handlers was sought during the measurement process. All measurements were taken with care to ensure accuracy and reliability.

### Measurement Procedure

The manual measurement method, by its nature, requires considerable attention, care, and time, and in certain circumstances, may also involve safety risks. Therefore, to enhance the accuracy of the data collected, each measurement was carefully recorded, and repeated verifications were performed by the same measures to ensure consistency. Given the challenges encountered during the measurement process, additional precautions were taken to improve the reliability of the data. Body measurements were conducted by a team of three individuals. During the procedure, one person was responsible for restraining the animal, while the other two performed the measurements (Fares, M. A., 2025). Due to the generally higher level of activity observed in female camels compared to males, additional safety measures were implemented when handling females. In some cases, assistance from caretakers was required to ensure that the animals remained stationary. During the measurements, some camels exhibited aggressive behaviors such as kicking or attempting to bite in response to contact with the measuring tape. In such instances, further precautions were taken to ensure the safety of both the animals and the research team, including calming measures to reduce stress and reactivity. The duration of each measurement session varied depending on the animal’s behavior, typically ranging between 5 and 10 minutes.

## Data Analysis

Within the scope of this study, selected body measurements of various Somali dromedary camel types located in Mogadishu were evaluated. The morphometric data collected in 2021 were digitized and subjected to statistical analysis. Microsoft Excel and the Statistical Package for the Social Sciences (SPSS) software were utilized for data processing and analysis. To determine significant differences among groups, one-way analysis of variance (ANOVA) was employed. Furthermore, Tukey’s post hoc multiple comparison test was conducted to identify statistically significant differences based on named type, sex, and named type × sex interaction. The results were thoroughly assessed to determine whether the body measurements of the camels exhibited significant variation as a function of named type and sex.

## Results

### Body Measurements

Based on the findings of the study, the body measurements of female individuals from different Somali dromedary camel types were comparatively evaluated (Table 2).

**Table 2.** Selected Body Measurements (cm) of Female Individuals Belonging to Different Camel Types.

| Measurement Place | SiifDacar (n=49) | Camel Types<br>Hoor (n=57) | Eyddimo (n=40) | Total (n=146) | p | Diff. |
| --- | --- | --- | --- | --- | --- | --- |
| Withers Height | 195.81±3.08<sup>a</sup> | 185.80±2.86<sup>b</sup> | 188.52±3.41 | 189.90±1.72 | **p<0.05** | a>b |
| Back Height | 229.87±7.56 | 214.47±7.00 | 207.78±8.36 | 217.81±5.70 | p>0.05 | - |
| Rump Height | 182.49±2.54<sup>a</sup> | 177.63±2.358<sup>b</sup> | 177.43±2.815<sup>c</sup> | 179.21±0.76 | **p<0.01** | a>b,c |
| Body Length | 154.06±1.15<sup>a</sup> | 148.74±1.06<sup>b</sup> | 149.27±1.268 | 150.67±0.70 | **p<0.01** | a>b |
| Chest Height | 132.39±3.56 | 141.11±3.28 | 141.13±3.94 | 138.18±2.64 | p>0.05 | - |
| Abdominal Height | 122.57±8.32 | 146.75±7.71 | 132.98±9.20 | 134.86±6.27 | p>0.05 | - |
| Shoulder Width | 33.94±0.40<sup>a</sup> | 31.95±0.38<sup>b</sup> | 31.73±0.45<sup>c</sup> | 32.55±0.27 | **p<0.01** | a>b,c |
| Seat Width | 24.94±0.35 | 24.05±0.32 | 24.45±0.383 | 24.44±0.20 | p>0.05 | - |

As shown in Table 2, statistically significant differences were found among specified types in terms of withers height, rump height, body length, and shoulder width in female camels (p < 0.05). On the other hand, no statistically significant differences were detected between the types for back height, chest height, abdominal height, and seat width (p > 0.05).

Regarding withers height, female camels of the SiifDacar type exhibited significantly greater values compared to those of the Hoor type (p < 0.05). This finding indicates that SiifDacar camels have a larger frame and a higher shoulder profile. Similarly, the rump height of female SiifDacar individuals was significantly higher than that of the Hoor and Eyddimo types (p < 0.01), suggesting a steeper and more elevated rump structure in the SiifDacar camels.

When body length was compared, SiifDacar females were found to be significantly longer than their Hoor counterparts (p < 0.01), indicating a more elongated body conformation. For shoulder width, individuals from the SiifDacar camels demonstrated significantly greater values compared to those of both the Hoor and Eyddimo camels (p < 0.01). Shoulder width is an important morphometric indicator in terms of muscle mass and body development, and a high value of this parameter indicates that Hoor camels have a more compact and strong body structure.

Based on the findings, female SiifDacar camels are thought to exhibit morphological characteristics closer to the dairy-type phenotype, with a higher withers height and a longer body structure. In contrast, Hoor and Eyddimo camels have lower withers height, but the Hoor camels exhibit a morphology more inclined to the meat-type profile due to their broader shoulder structure.

Additionally, Table 3 presents selected morphometric measurements of male individuals from different Somali dromedary camel types.

**Table 3.**
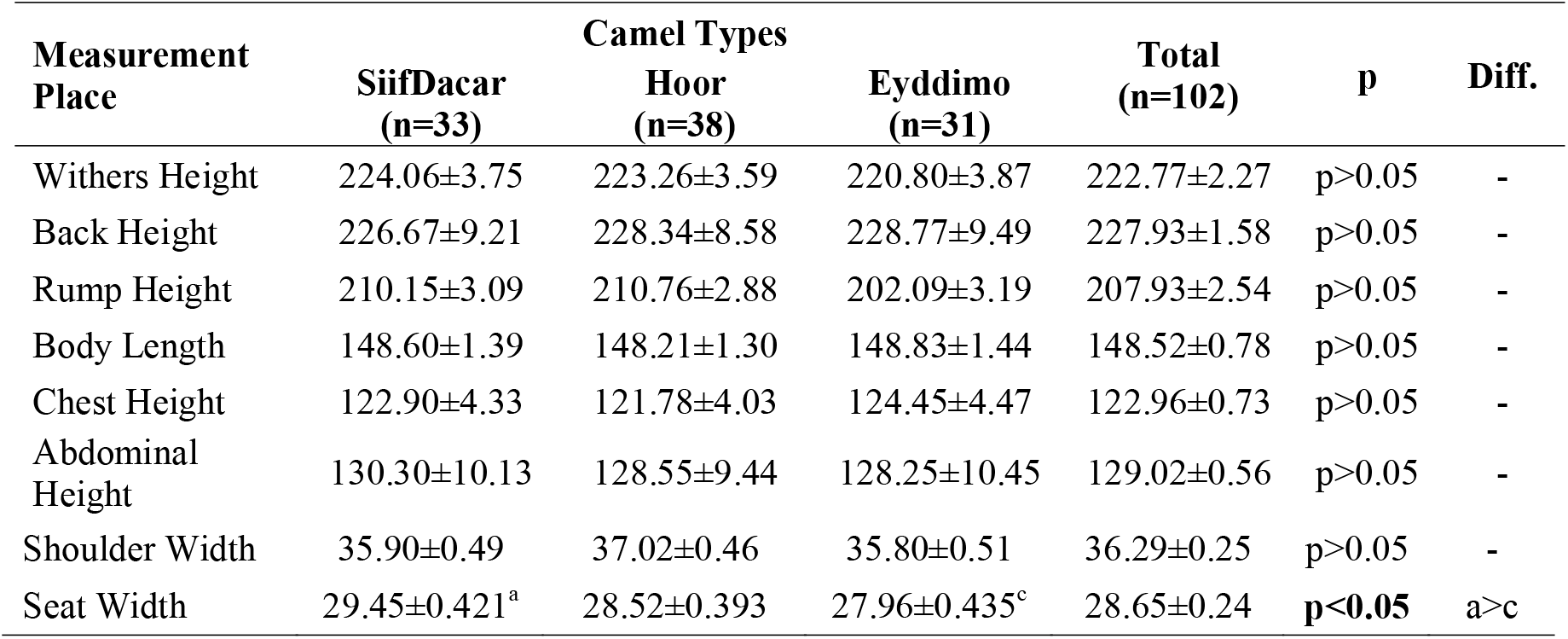
Selected Body Measurements (cm) of Male Individuals Belonging to Different Camel Types.

The analyses revealed no statistically significant differences between reported camel types for withers height, back height, rump height, body length, chest height, abdominal height, and shoulder width (p > 0.05). However, a statistically significant difference was found for seat width (p < 0.05), with male individuals of the SiifDacar camels exhibiting a broader rump structure compared to those of the Eyddimo camels.

The findings indicate that, although the overall body structure of male Somali dromedary camels is similar across types, there are distinct differences in seat width. Given that seat width is related to feeding efficiency and muscle development, the SiifDacar camels may exhibit a more advantageous morphological trait in this regard.

### Sex and Type Comparisons

In this study, some body measurements of different Somali dromedary camel types within the age range of 6-9 years were examined, and changes related to sex and type × sex interactions were statistically evaluated (Table 4).

**Table 4.**
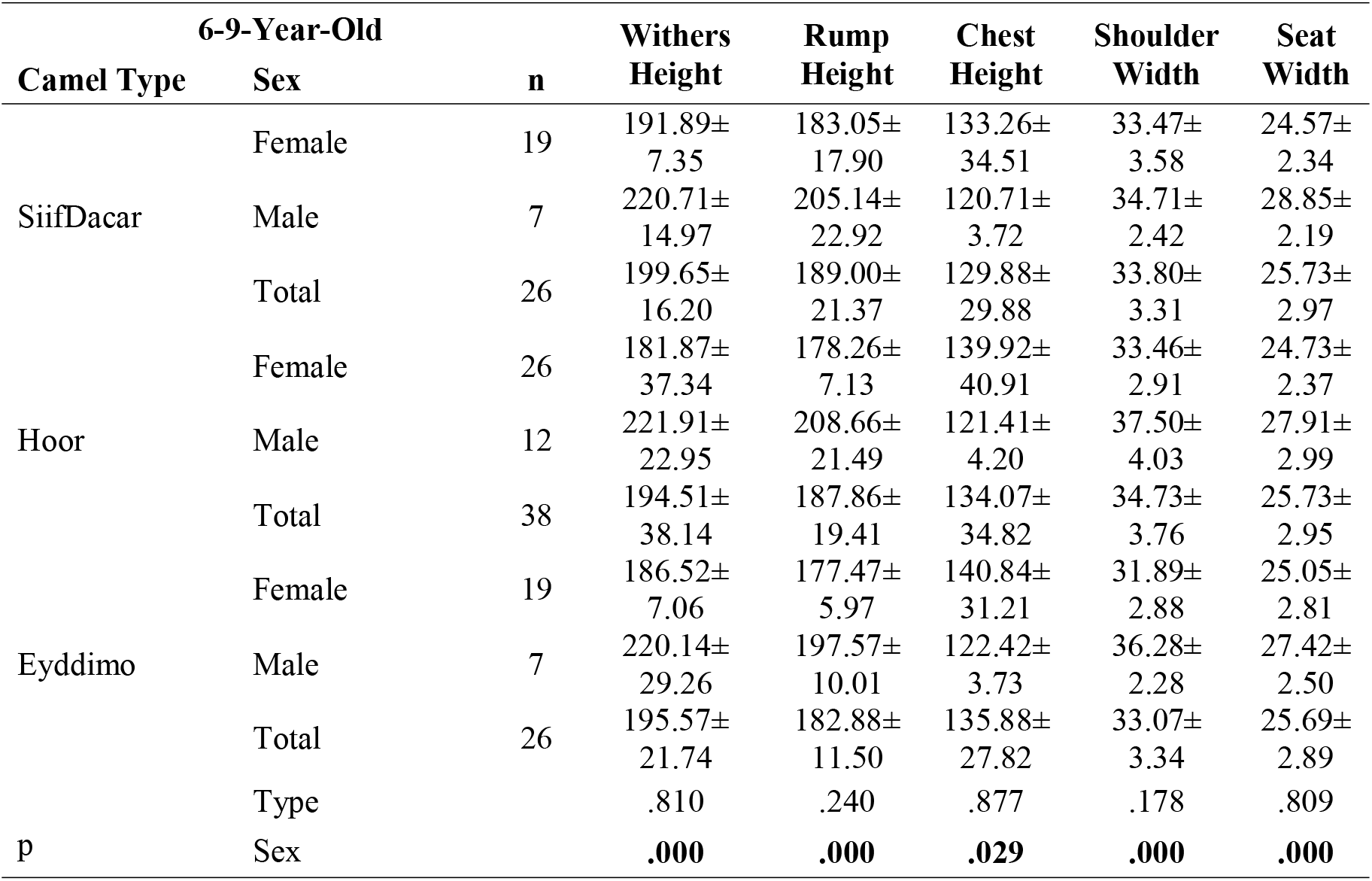
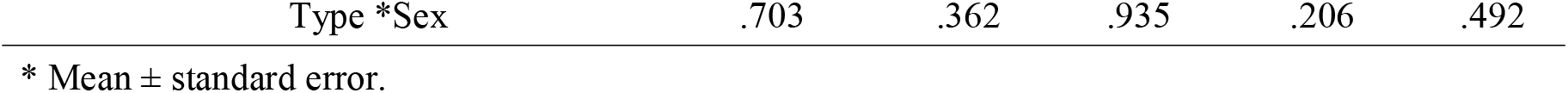
Morphometric Characteristics of 6–9-Year-Old Individuals in Different Camel Types Based on Sex and Type × Sex Interaction.

As seen in Table 4, the analysis results reveal that the parameters of withers height, rump height, chest height, shoulder width, and seat width did not show statistically significant differences between the reported camel types (p > 0.05). However, significant differences based on sex were observed (p < 0.05). Furthermore, it was determined that there was no statistically significant effect of the interaction between type and sex on the measured characteristics (p > 0.05).

When comparing the sexes, it was observed that male camels had significantly larger values for withers height, rump height, and shoulder width compared to females. Particularly, male camels of the Hoor camels had higher shoulder width values compared to other groups, indicating that this camel type may have an advantage in terms of meat production. For chest height, a significant difference was found between sexes, with females having higher values compared to males (p < 0.05). This finding suggests that there may be regional differences in the body structure of female individuals. Therefore, Table 5 presents the morphometric characteristics of different Somali dromedary camel types for individuals aged 10-13 years, based on sex and type × sex interaction.

**Table 5.**
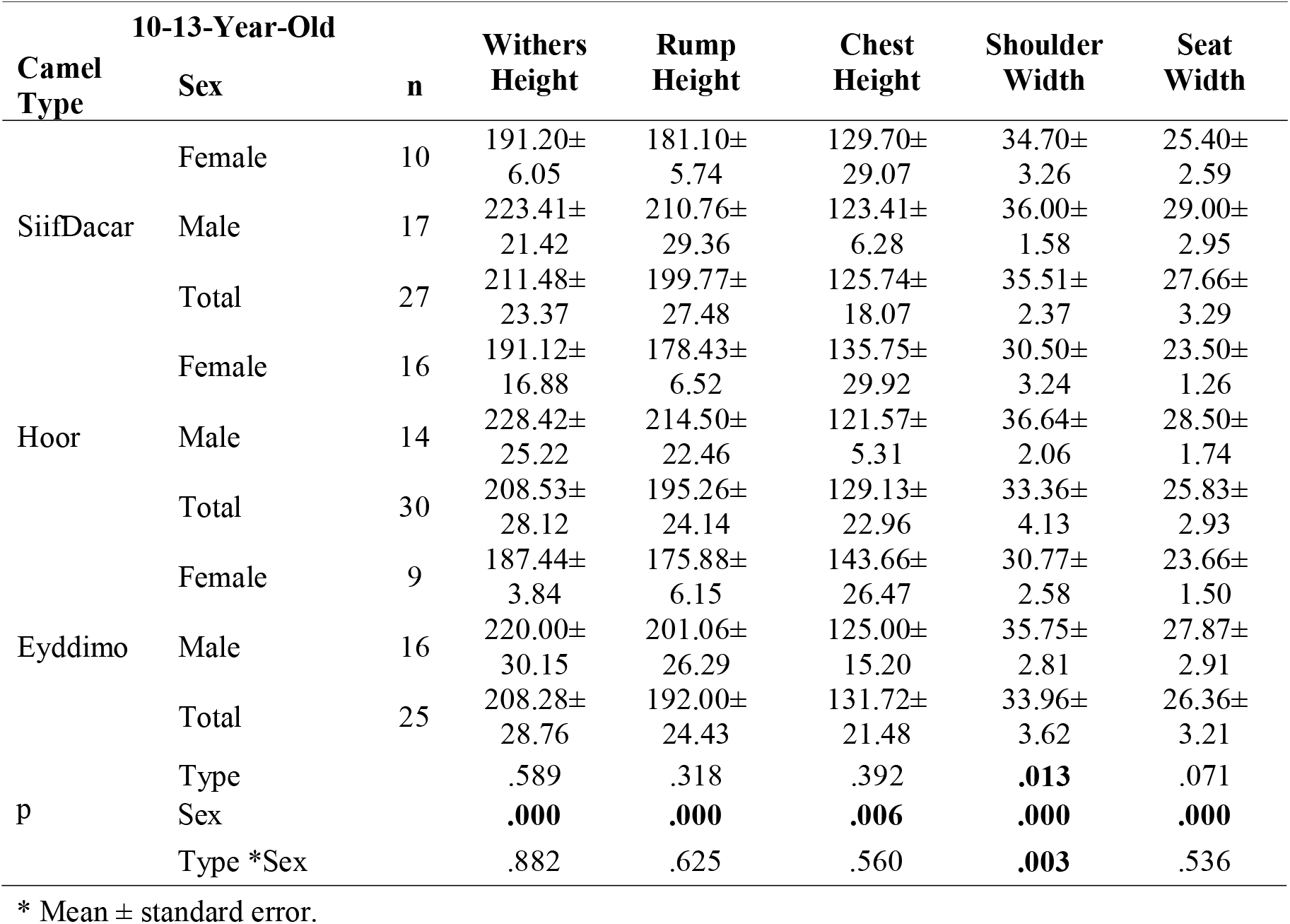
Morphometric Characteristics of Individuals Aged 10-13 Years from Different Camel Types Based on Sex and Type × Sex Interaction.

As seen in Table 5, there was no statistically significant difference between the reported types in terms of withers height, rump height, chest height, shoulder width, and seat width for camels aged 10-13 years (p>0.05). However, a statistically significant difference was found in terms of shoulder width (p=0.013). Comparisons based on sex revealed that, for all parameters, male camels had significantly larger values than females (p<0.05). Additionally, a statistically significant interaction between camel type and sex was found for shoulder width (p=0.003).

When sex differences were evaluated, it was observed that male camels had noticeably larger values than females in terms of withers height, rump height, shoulder width, and seat width. This can be explained by the fact that male camels tend to have a larger morphological structure due to genetic and hormonal factors.

Male camels of the Hoor and SiifDacar camels had higher values for withers and rump height compared to other groups. For chest height, female camels were found to have higher values than males (p<0.05), which suggests that female camels may have a broader chest area, and this difference could be related to physiological or structural adaptations.

The presence of a significant difference in shoulder width between the reported camel types indicates that this morphometric feature may be influenced by camel type-specific genetic and environmental factors. Specifically, male camels of the Hoor camels had the highest shoulder width values, suggesting that this type may have a stronger muscular structure.

Table 6 shows some body measurements of different Somali dromedary camel types at the age of 14 years and older, and the changes due to gender and type*gender interaction.

**Table 6.**
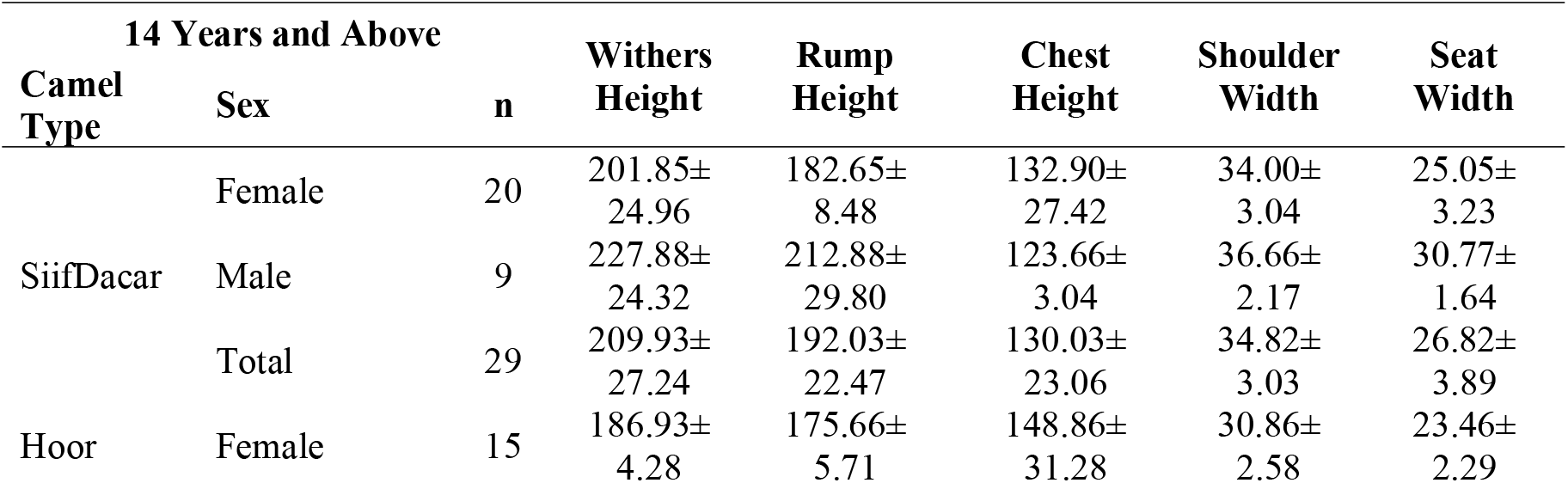
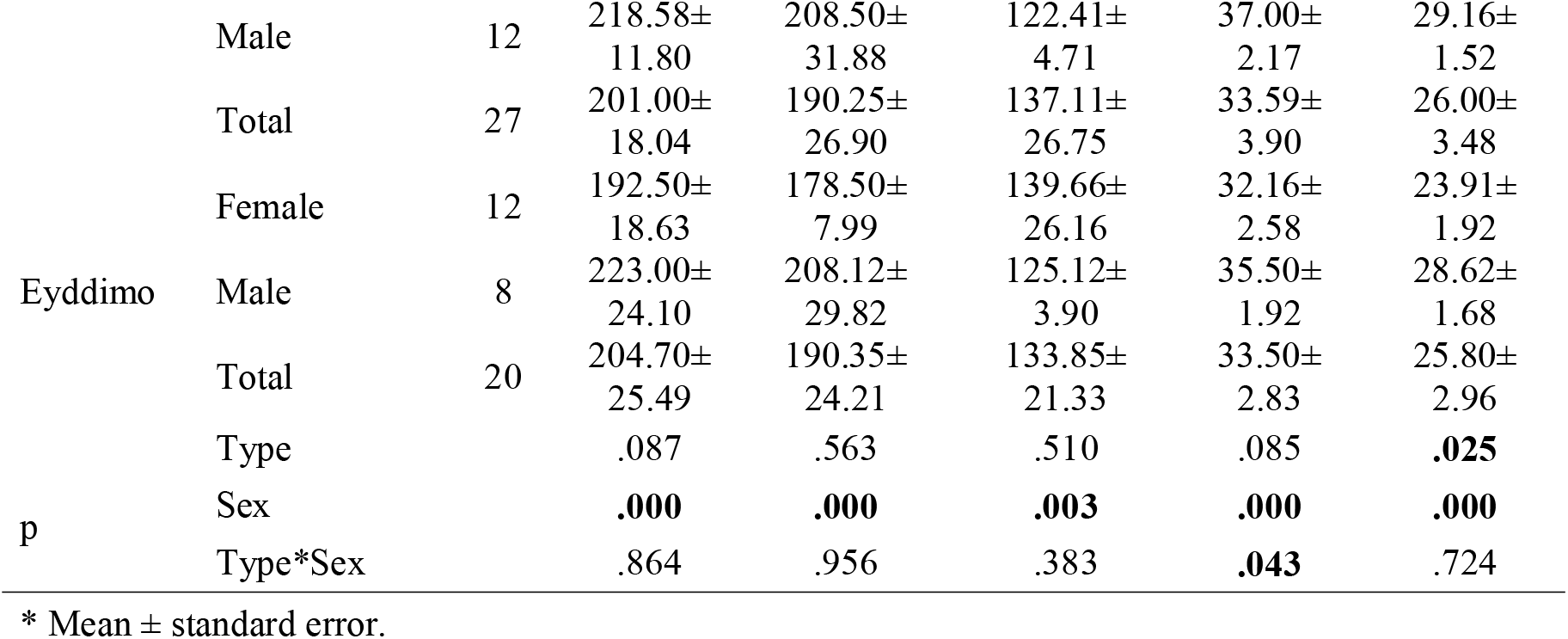
Morphometric Characteristics of Individuals Aged 14 and Above from Different Camel Types Based on Sex and Type × Sex Interaction.

The research findings indicate that there were generally no statistically significant differences among the reported camel types in terms of withers height, rump height, chest height, shoulder width, and seat width (p>0.05). However, a statistically significant difference was observed among different types in the parameter of seat width (p=0.025). When evaluated based on sex, male individuals were found to have significantly higher values across all body measurements compared to females (p<0.05). Moreover, the interaction between camel type and sex was determined to be statistically significant for shoulder width (p=0.043).

When sex-related differences were examined, it was observed that males had notably higher values for withers height, rump height, shoulder width, and seat width. The greater withers and rump heights observed in males suggest that they may possess a larger and more robust skeletal structure compared to females. Notably, male individuals belonging to the Hoor and SiifDacar camels exhibited the highest values for withers and rump height. In contrast, comparisons of chest height revealed that female individuals had significantly higher values than males (p=0.003). This difference may be attributed to the broader thoracic structure in females and is likely linked to physiological adaptations. Regarding shoulder width, males were found to have significantly broader shoulders than females (p<0.05). The fact that males of the Hoor camels recorded the highest shoulder width values suggests that this camel type may exhibit distinct structural characteristics compared to others. Additionally, the statistically significant interaction between camel type and sex in shoulder width (p=0.043) indicates that this trait is influenced by both genetic factors and environmental conditions.

### Correlation Among Body Measurements

Table 7 presents the correlation findings among body measurements of the camels.

**Table 7.** Correlation Findings Between Body Measurements.

|  |  | (1) | (2) | (3) | (4) | (5) | (6) | (7) | (8) |
| --- | --- | --- | --- | --- | --- | --- | --- | --- | --- |
| **Withers Height (1)** | r | 1 |  |  |  |  |  |  |  |
|  | p |  |  |  |  |  |  |  |  |
| **Back Height (2)** | r | 0.051 | 1 |  |  |  |  |  |  |
|  | p | 0.427 |  |  |  |  |  |  |  |
| **Rump Height (3)** | r | 0.449 | 0.068 | 1 |  |  |  |  |  |
|  | p | **0.000** | 0.289 |  |  |  |  |  |  |
| **Body Length (4)** | r | 0.085 | -0.070 | 0.200 | 1 |  |  |  |  |
|  | p | 0.184 | 0.275 | **0.002** |  |  |  |  |  |
| **Chest Height (5)** | r | -0.224 | 0.119 | -0.204 | -0.015 | 1 |  |  |  |
|  | p | **0.000** | 0.062 | **0.001** | 0.819 |  |  |  |  |
| **Abdominal Height (6)** | r | -0.087 | -0.081 | -0.073 | 0.022 | -0.068 | 1 |  |  |
|  | p | 0.173 | 0.201 | 0.249 | 0.713 | 0.283 |  |  |  |
| **Shoulder Width (7)** | r | 0.421 | 0.093 | 0.383 | 0.071 | -0.141 | -0.098 | 1 |  |
|  | p | **0.000** | 0.142 | **0.000** | 0.267 | **0.026** | 0.126 |  |  |
| **Seat Width (8)** | r | 0.500 | 0.040 | 0.528 | 0.015 | -0.175 | -0.117 | 0.569 | 1 |
|  | p | **0.000** | 0.529 | **0.000** | 0.817 | **0.006** | 0.066 | **0.000** |  |

As shown in Table 7, statistically significant relationships were identified between certain morphometric traits of the animals analyzed.

Strong positive correlations were found between withers height and rump height (r = 0.449, p < 0.001), shoulder width (r = 0.421, p < 0.001), and seat width (r = 0.500, p < 0.001). These results indicate that as wither height increases, rump height, shoulder width, and seat width also tend to increase.

Significant positive associations were also observed between rump height and body length (r = 0.200, p = 0.002), seat width (r = 0.528, p < 0.001), and shoulder width (r = 0.383, p < 0.001). This suggests that individuals with greater rump height generally exhibit broader rump and shoulder structures. Additionally, individuals with longer body length tend to have higher rump height.

A moderate positive correlation was detected between shoulder width and seat width (r = 0.569, p < 0.001), implying that camels with broader shoulders often have greater seat width as well.

Negative correlations were identified between chest height and withers height (r = -0.224, p < 0.001), as well as rump height (r = -0.204, p = 0.001). These findings suggest a tendency for chest height to decrease as withers and rump heights increase.

Furthermore, chest height was also negatively correlated with shoulder width (r = -0.141, p = 0.026), indicating that individuals with broader shoulders may have relatively shallow chest height. A similar negative relationship was observed between chest height and seat width (r = -0.175, p = 0.006), suggesting that those with greater chest height might exhibit comparatively narrower seat width.

Therefore, the correlation analysis revealed significant positive relationships among key body measurements, such as withers height, rump height, and shoulder width. On the other hand, measurements like chest height and abdominal height showed inverse relationships with several other traits. These findings provide valuable insights into the morphometric structure of Somali dromedary camels and contribute to the identification of optimal morphological characteristics for breeding purposes.

## Discussion

This study aimed to determine and statistically evaluate specific body measurements and their interrelationships among three distinct types of Somali dromedary camels, SiifDacar, Hoor, and Eyddimo, raised in farms located in the Dharkenley, Dayniile, and Hodan districts of Mogadishu. A total of 248 camels, including 102 males and 146 females, were subjected to morphometric measurements.

According to the findings, significant differences were observed among the female camels of different types in terms of withers height, rump height, body length, and shoulder width. A similar study has been done by Isako and Kimindu (2019). The SiifDacar camels exhibited greater withers and rump height compared to the Hoor camels. Although no significant difference in body length was found between the Hoor and Eyddimo camels, the SiifDacar camels displayed a longer body structure than both. Similarly, while shoulder width did not differ between Hoor and Eyddimo, SiifDacar females had a notably broader shoulder structure. Overall, the SiifDacar camels showed larger body dimensions in comparison to the other two. These characteristics, taller stature and longer body in SiifDacar females, suggest that this type may possess morphological features more aligned with dairy-type camels. In contrast, the broader shoulder structure of the Hoor camels may indicate a body conformation more suited to meat production, similar to a study that has been shown (Ishag *et al*., 2011).

In male camels, a statistically significant difference was observed among the reported types only for seat width. SiifDacar males exhibited the broadest rumps, while the Eyddimo camels had the narrowest, with Hoor falling in between. Although the general body structure of the males appeared relatively similar across types, the variation in seat width stands out. Given that seat width is associated with fattening performance and muscle development, the SiifDacar camels may have a morphological advantage in this regard (Jafar, S. (2018).

In age-based evaluations, no significant differences among camel types were found in the 6– 9-year-old age group for withers height, rump height, chest height, shoulder width, or seat width. Instead, sex emerged as a determining factor: males had greater withers and rump height, while females displayed greater chest height. Similarly, in the 10–13-year age group, sex-related differences were again prominent, with males exhibiting generally larger measurements than females (Kebede *et al*., 2022). However, shoulder width varied significantly based on both sex and camel type. Among camels aged 14 and above, parameters such as withers height, back height, and chest height were predominantly influenced by sex, with male camels showing more robust structures (Legesse *et al*. 2018). Notably, the statistically significant differences in seat width among camel types in this age group suggest that this trait may be influenced by the reported type-specific genetic variation. Identifying such structural distinctions could contribute to improving breeding strategies and refining selection programs (Leyland *et al*, 2014).

This study also explored the correlations among body measurements, including withers height, back height, rump height, body length, chest height, abdominal height, shoulder width, and seat width in both male and female camels. Moderate positive correlations were found between withers height and rump height (r = 0.449), as well as between withers height and shoulder width (r = 0.421). Relationships between rump height and body length (r = 0.200) and between rump height and seat width (r = 0.528) were also positively correlated, though to a lesser degree. Weak negative correlations were detected between chest height and withers height (r = -0.224), as well as between chest height and rump height (r = -0.204). A weak positive correlation was observed between shoulder width and rump height (r = 0.383), while shoulder width and chest height (r = -0.141) showed a very weak negative relationship (Li *et al*., 2017). A moderate positive correlation was found between shoulder width and seat width (r = 0.569).

This research determined that, on average, the Mogadishu camels had a withers height of 189.90 cm, back height of 217.81 cm, rump height of 179.21 cm, body length of 150.67 cm, chest height of 138.18 cm, abdominal height of 134.86 cm, shoulder width of 32.55 cm, and seat width of 24.44 cm. When compared to the findings of (Li *et al*., 2017; Çağlı, 2019), who studied camels in Türkiye, it was observed that the Mogadishu camels had higher withers and rump height but narrower shoulders. Similarly, Yılmaz et al. 2013; Meghelli *et.al*. 2020) reported a withers height of 161.70 cm in Turkish wrestling camels, a value lower than that observed in the present study. Ishag et al. (2011) found an average withers height of 189.42 cm among female camels in Sudan, which aligns closely with the values recorded in Mogadishu. Razig et al. 2011; Tandoh, *et.al*., 2018; Yakubu, *et.al*., 2022. In their study of camels from Pakistan and Afghanistan (Muigai *et.al*., 2016; Mushtaq *et.al*., 2025). reported a withers height of 164 cm and a rump height of 139 cm, both of which are lower than those found in this research.

In the study by Raziq et.al. (2011) and Çağlı (2019), strong positive correlations were reported between withers height and rump height in both male and female camels. High positive correlations were also observed between body length, back height, shoulder width, and seat width (Yılmaz *et.al*., 2013; Yosef *et.al*., 2014). The correlation findings in the present study are consistent with those reported by Çağlı (2019). In this context, determining the morphometric characteristics of different camel types serves as a valuable data source for the development of breeding programs and improvement of genetic selection processes.

## Conclusion

This study was carried out to determine the morphometric characteristics of the camels named as different dromedary camels raised in Mogadishu according to various age and gender groups, and in light of the obtained data, it was determined that there were significant differences between the named camels. Therefore, these differently named camels are thought to be separate breeds. The SiifDacar camels were generally found to possess larger body measurements, whereas the Hoor and Eyddimo camels had comparatively smaller body structures. Sex was also identified as a significant factor influencing body dimensions, with males typically displaying more robust physiques than females. These results have the potential to contribute to the development of breeding strategies, performance evaluations, and breed registration efforts.

The data obtained from this research provide valuable insights into camel husbandry practices in Somalia and aim to fill a scientific gap in the field. Further detailed studies are needed to assess milk and meat productivity in relation to both genetic and environmental factors. Existing literature reports a wide variation in milk yield among camels (Çağlı, 2019), highlighting the importance of investigating the genetic and environmental determinants of milk production in camels raised in Mogadishu.

This research also represents an important starting point for understanding the genetic similarities among different camel types and breeds. Studies have shown that the body measurements of camels raised in Türkiye differ from those of Mogadishu camels (Yılmaz et al., 2013; Çağlı, 2019). It remains to be clarified whether these differences arise from genetic or environmental influences. Therefore, conducting genetic comparisons among camel populations from diverse regions such as Türkiye, Sudan, Pakistan, and Afghanistan will be vital for identifying inter-population similarities and differences.

Furthermore, it is recommended that future research explores in greater detail the relationship between the body measurements of Mogadishu camel types and economically important traits such as milk and meat productivity. Selection programs based on these data could enhance the efficiency of camel husbandry in the region. In this context, future studies should focus on evaluating the production potential of these camel types and developing sustainable breeding strategies. The findings of this study are expected to contribute to both local and international research efforts related to camel breeding and management.

## ACKNOWLEDGEMENT

We would like to express our sincere gratitude to Selçuk University for providing academic support and a conducive research environment. We also extend our appreciation to the Department of Animal Production and Development at the Ministry of Livestock, Forestry and Range, Somalia, for facilitating fieldwork and access to study animals in Mogadishu.

Special thanks are due to Prof. Dr. Alper Yılmaz for his invaluable guidance, supervision, and constructive comments throughout the study. We also acknowledge the support of local livestock owners and assistants who contributed to data collection.

## ETHICS STATEMENT

This study was conducted in accordance with accepted standards for animal research and welfare. All procedures involving animals were performed following institutional and international guidelines for the care and use of animals in research.

Ethical approval for this study was obtained from Selçuk University Animal Experiments Ethics Committee (Approval No. SÜVDAMEK2021/77). In addition, permission to conduct fieldwork and collect data was granted by the Ministry of Livestock, Forestry, and Range, Somalia. Therefore, Verbal consent was obtained from camel owners before inclusion of their animals in the study.

## Author Contributions

Hassan Hussein Abdi contributed to the study conception, data collection, analysis, and drafting of the manuscript. Prof. Dr. Alper Yılmaz supervised the research, contributed to the study design, and critically revised the manuscript. Abdirahman Barre contributed to data analysis, interpretation of results, and manuscript revision. Prof. Dr. Feaz Jesse Abdullah contributed to methodological guidance, data interpretation, and critical review of the manuscript. All authors read and approved the final manuscript.

## GENERATIVE AI STATEMENT

The author(s) declare that no Generative AI was used in the creation of this manuscript. Any alternative text (alt text) provided alongside figures in this article has been generated by Frontiers with the support of artificial intelligence, and reasonable efforts have been made to ensure accuracy, including review by the authors wherever possible. If you identify any issues, please contact us.

## Availability of Data and Materials

The datasets used and/or analyzed during the current study are available from the corresponding author on reasonable request.

